# *In vivo* augmentation of a complex gut bacterial community

**DOI:** 10.1101/2021.06.15.448620

**Authors:** Alice G. Cheng, Po-Yi Ho, Sunit Jain, Xiandong Meng, Min Wang, Feiqiao Brian Yu, Mikhail Iakiviak, Ariel R. Brumbaugh, Kazuki Nagashima, Aishan Zhao, Advait Patil, Katayoon Atabakhsh, Allison Weakley, Jia Yan, Steven Higginbottom, Norma Neff, Justin L. Sonnenburg, Kerwyn Casey Huang, Michael A. Fischbach

**Affiliations:** Department of Gastroenterology, Stanford School of Medicine, Stanford, CA 94305, USA; Department of Bioengineering, Stanford University, Stanford, CA 94305, USA; Chan Zuckerberg Biohub, San Francisco, CA 94158, USA; Federation Bio, South San Francisco, CA 94080; Department of Microbiology and Immunology, Stanford University School of Medicine, Stanford University, Stanford, CA 94305, USA; ChEM-H Institute, Stanford University, Stanford, CA 94305, USA

## Abstract

Efforts to model the human gut microbiome in mice have led to important insights into the mechanisms of host-microbe interactions. However, the model communities studied to date have been defined or complex but not both, limiting their utility. In accompanying work, we constructed a complex synthetic community (104 strains, hCom1) containing the most common taxa in the human gut microbiome. Here, we used an iterative experimental process to improve hCom1 by filling open metabolic and/or anatomical niches. When we colonized germ-free mice with hCom1 and then challenged it with a human fecal sample, the consortium exhibited surprising stability; 89% of the cells and 58% of the taxa derive from the original community, and the pre- and post-challenge communities share a similar overall structure. We used these data to construct a second version of the community, adding 22 strains that engrafted following fecal challenge and omitting 7 that dropped out (119 strains, hCom2). In gnotobiotic mice, hCom2 exhibited increased stability to fecal challenge and robust colonization resistance against pathogenic *Escherichia coli*. Mice colonized by hCom2 versus human feces are similar in terms of microbiota-derived metabolites, immune cell profile, and bacterial density in the gut, suggesting that this consortium is a prototype of a model system for the human gut microbiome.

## INTRODUCTION

Experiments in which a microbial community is transplanted into germ-free mice have opened the door to studies of mechanism and causality in the microbiome. These efforts fall into two categories based on the nature of the transplanted community: studies involving transplantation of complete, undefined communities (i.e., fecal samples) have shown that the microbiome plays a role in a variety of host phenotypes including the response to cancer immunotherapy (Gopalakrishnan et al. 2018; Matson et al. 2018; Routy et al. 2018), caloric harvest (Ridaura et al. 2013), colonization resistance to enteric pathogens (Buffie et al. 2015), and neural development (Sharon et al. 2019; Buffington et al. 2021). While illuminating, a limitation of this format is that it is difficult to ‘fractionate’ an undefined community, making it challenging to discover which strains are involved in a phenotype of interest.

As an alternative, germ-free mice have been colonized with communities that are incomplete but defined (i.e., a synthetic community). These studies have revealed mechanistic insights into immune modulation, glycan consumption, and other complex phenotypes driven by the microbiome (Wymore Brand et al. 2015; Patnode et al. 2019; Faith et al. 2014). Synthetic communities enable precise manipulations such as strain dropouts and gene knockouts. However, the communities used are typically of low complexity, limiting their ability to model the biology of a native-scale microbiome.

An ideal model system for the gut microbiome would capture the advantages of both approaches. To this end, we sought to create a community that is completely defined, enabling precise manipulations, but complex enough to exhibit emergent features of a complete community (e.g., stability upon transplantation, colonization resistance). We started by colonizing germ-free mice with a defined 104-member community (hCom1) designed in an accompanying effort (Cheng et al. 2021), showing that it adopts a stable, highly reproducible configuration in which its constituent strains span seven orders of magnitude of relative abundance. We augmented the community, filling open niches using an iterative ecology-based process, and we show that the enlarged community (hCom2) is more resilient to perturbation and resistant to pathogen colonization. Mice colonized by hCom2 are phenotypically similar to mice harboring an undefined human fecal sample, suggesting that our consortium and augmentation process lay the foundation for developing defined models of the human gut microbiome.

## RESULTS

### Attributes of a complex defined community in gnotobiotic mice

In accompanying work (Cheng et al. 2021), we designed a 104-member community (hCom1) that includes many of the most common bacterial taxa from the human gut microbiome. As a starting point for this study, we colonized germ-free Swiss-Webster (SW) mice with hCom1 (**Figure 1A**), which we prepared by propagating each strain individually and mixing OD-normalized cultures (**Methods**). We sampled fecal pellets from the mice weekly for eight weeks, enumerated community composition in the inoculum and each fecal sample by metagenomic sequencing, and performed read analysis with a highly sensitive and specific read mapping pipeline, NinjaMap (Cheng et al. 2021). Taking advantage of the fact that each organism in hCom1 is sequenced, NinjaMap translates read-counting statistics into community composition information even for low-abundance organisms (<10^−6^). Our analysis yielded three main conclusions:

**Figure 1:**
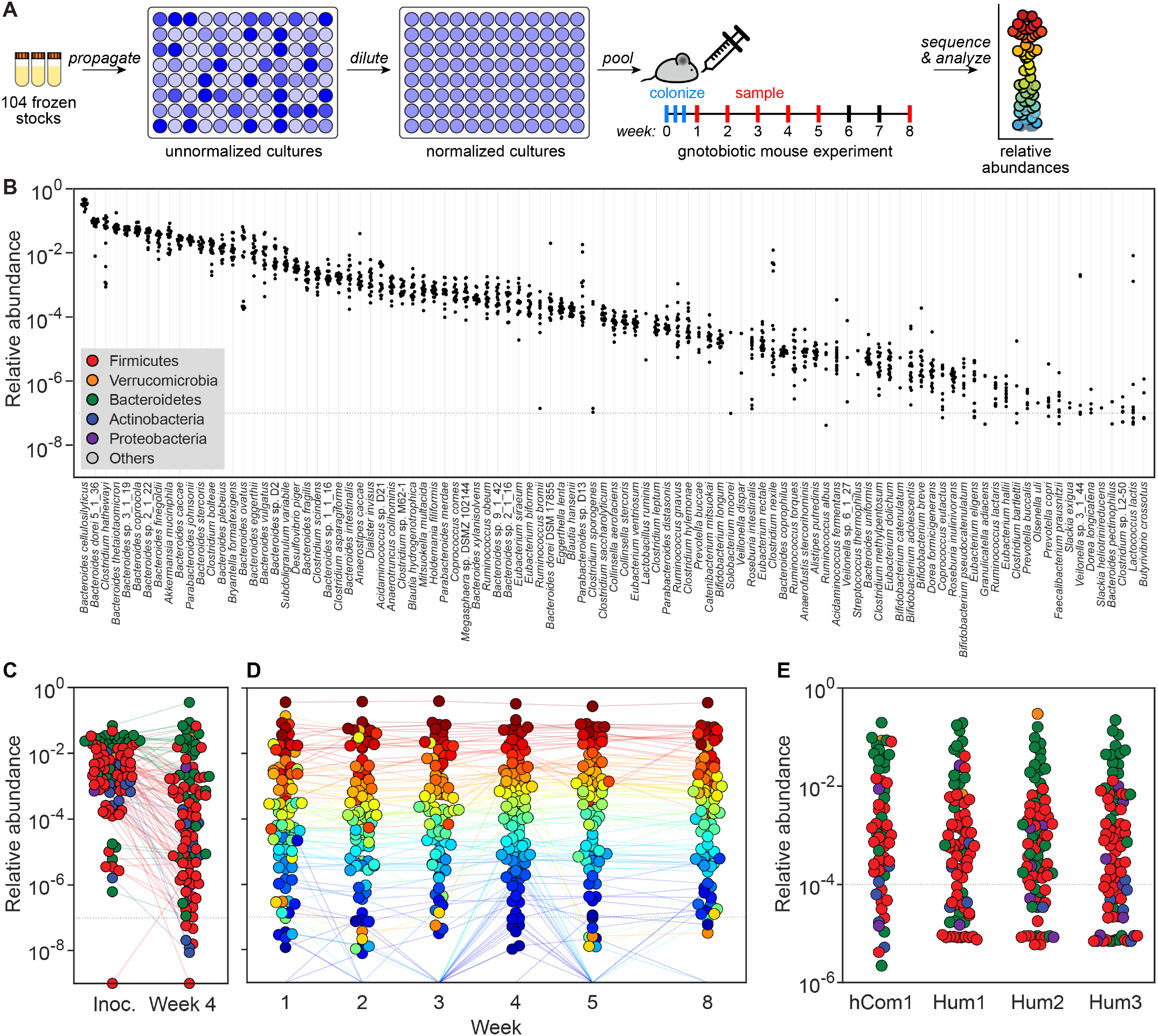
Colonizing germ-free mice with a complex gut bacterial community. (**A**) Schematic of the experiment. Frozen stocks of the 104 strains were used to inoculate cultures that were grown for 24 h, diluted to similar optical densities (to the extent possible), and pooled. The mixed culture was used to colonize germ-free Swiss-Webster (SW) mice by oral gavage. Fecal samples were collected at weeks 1-5 and week 8, subjected to metagenomic sequencing, and analyzed by NinjaMap to measure the composition of the community at each timepoint. (**B**) Relative abundances for most strains are tightly distributed. Each column depicts the relative abundance of an individual strain across all samples at week 4. (**C**) Averaged relative abundances of the inoculum versus the communities at week 4. Strains in the community span >6 orders of magnitude of relative abundance when colonizing the mouse gut. Dots are colored by phylum according to the legend in panel B. (**D**) hCom1 reaches a stable configuration by week 2. Each dot is an individual strain; the collection of dots in a column represents the community at a single timepoint averaged over 5 mice co-housed in a cage. Strains are colored according to their rank-order relative abundance at week 4. (**E**) The architecture of the community resembles that of a complete, undefined human fecal consortium. Averaged relative abundances of MIDAS bins—a rough proxy for species—are shown for hCom1-colonized mice versus mice colonized by stool from one of three healthy human donors (Hum1-3). Dots are colored by phylum according to the legend in panel B. In each case, the distribution of log relative abundances was centered at ∼0.01% and the median relative abundance of Bacteroidetes was higher than that of the Firmicutes.

First, almost all strains in the inoculum colonize the mouse gut (**Figure 1B-C**). We confirmed the presence of 103/104 strains in the inoculum; of these, 101 strains were detected in the mice at least once. While strain relative abundances spanned >7 orders of magnitude, nearly all strains exhibited low variation across 20 mice in four cages, with coefficient of variation (CV, standard deviation/mean) <0.4.

Second, the community quickly reaches a stable configuration (**Figure 1D**). Averaged across mice, relative abundances remained mostly constant starting two weeks after colonization, with Pearson’s correlation coefficient >0.95 at each time point with respect to the composition in week 8. After the first week, relative abundances stayed within a narrow range for the duration of the experiment (mean CV<0.2 across the 96 strains that remained above the limit of detection). Large shifts in relative abundance were rare: only 27/312 (8.7%) week-to-week strain-level changes were >10-fold.

Third, the architecture of the community resembles that of a complete, undefined human fecal consortium (**Figure 1E**). We colonized germ-free mice with three human fecal samples (hereafter, ‘humanized’) and compared their community compositions to those of mice colonized with hCom1. For this analysis we used MIDAS (Nayfach et al. 2016), an enumeration tool that—unlike NinjaMap—does not require prior knowledge of the constituent strains. MIDAS and NinjaMap reported highly concordant relative abundance profiles using sequencing reads from hCom1-colonized mice, although—as expected—MIDAS was less sensitive since it utilizes only 1% of sequencing reads (**Figure S1**). We used MIDAS for subsequent analyses of samples that are partially or completely undefined.

The gut communities of hCom1-colonized and humanized mice were similar in three ways: (*i*) Relative abundances spanned at least five orders of magnitude, with some strains consistently colonizing at >10% and others at <0.001%. (*ii*) The distribution of log relative abundances was centered at ∼0.01%, indicating that the majority of strains in the community would be missed by enumeration tools that have a limit of detection of 0.1%. (*iii*) Phylum-level distributions are similar: the median relative abundance of Bacteroidetes was higher than that of the Firmicutes (2.4×10^−2^ versus 7.7×10^−4^ in hCom1-colonized mice, with total abundances of 0.78 and 0.12 for Bacteroidetes and Firmicutes, respectively; 8.1×10^−3^ versus 2.7×10^−4^ in a representative humanized mouse, with total abundances of 0.84 and 0.12). In both cases, Actinobacteria were present at low relative abundance (∼10^−4^). Thus, hCom1 takes on a similar architecture to a human fecal community in the mouse gut.

### An ecology-based process to fill open niches in the community

Although hCom1 is complex and phylogenetically representative of the human gut microbiome, it is not as complex or phylogenetically rich as a human fecal community (**Figure 1E**); indeed, the criteria that dictated its membership were not designed to ensure completeness by any functional or ecological criteria (Cheng et al. 2021). To create a defined community that better approximates the gut microbiome, we sought to augment hCom1 by increasing the number of niches it fills in the gastrointestinal tract. We designed an experimental strategy to fill open niches using strains from a complete consortium (**Figure 2A**). Taking advantage of colonization resistance (Buffie and Pamer 2013; Lawley and Walker 2013), an ecological phenomenon in which resident organisms exclude invading strains from occupied niches, we started by colonizing germ-free mice for four weeks with hCom1, presumably filling the metabolic and anatomical niches in which its strains reside. We then challenged the mice with one of three undefined fecal samples, reasoning that invading strains that would otherwise occupy a niche already filled by hCom1 would be excluded, whereas invading strains whose niche was unfilled would be able to cohabit with hCom1. After four additional weeks, we used metagenomic sequencing to analyze community composition from fecal pellets.

**Figure 2:**
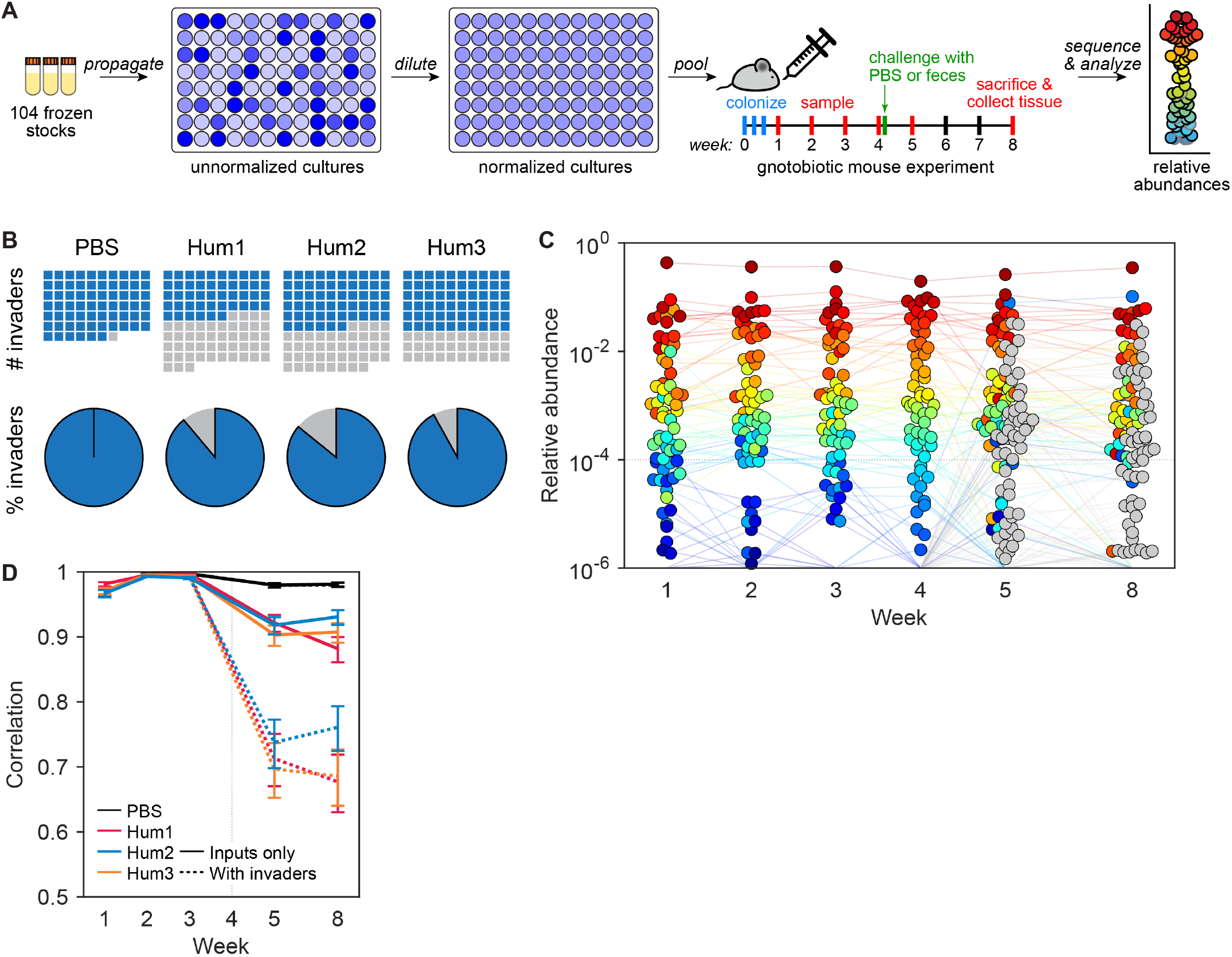
Challenging hCom1 with human fecal communities to identify strains that fill open niches. (**A**) Schematic of the experiment. Mice were colonized by hCom1 and housed for four weeks, presumably filling the metabolic and anatomical niches accessible to the strains in the community. At the beginning of week 5, the mice were challenged with one of three fecal communities from a healthy human donor or with PBS as a control; we reasoned that fecal strains that would otherwise occupy a niche already filled by hCom1 would be excluded, whereas fecal strains whose niche was unfilled would be able to cohabit with hCom1. After four additional weeks, we used metagenomic sequencing coupled with MIDAS to analyze community composition from fecal pellets collected at weeks 1-5 and 8. (**B**) hCom1 was broadly but not completely resistant to fecal challenge. Top row: Blue squares in the waffle plots indicate strains that derive from hCom1, and gray squares represent strains from the fecal communities. Bottom row: pie charts representing the percentage of MIDAS bins, a rough proxy for species-level taxa, that derive from hCom1 versus the fecal communities. An average of 89% of the genome copies from week 8—and 58% of the MIDAS bins, a rough proxy for species—derived from hCom1. The remaining 11% of the genome copies, and 42% of the MIDAS bins, represent new species that joined hCom1 from one of the fecal samples. (**C**) Despite the addition of new strains, the architecture of the community remained intact. Each dot is an individual strain; the collection of dots in a column represents the community at a single timepoint averaged over the 5 co-housed mice that were challenged with fecal community Hum1. Strains are colored according to their rank-order relative abundance at week 4. Gray circles represent invading species, defined as any species not present in weeks 1-4 in the group of mice shown. (**D**) The relative abundances of the hCom1-derived species present post-challenge were highly correlated with their pre-challenge levels. Pearson’s correlation coefficient with respect to the average relative abundance in weeks 3 and 4 are shown for the PBS control and 3 fecal community challenges, averaged across mice that received the same challenge. Correlation coefficients are shown for the 104 hCom1 species (solid lines) and for all species including invaders (dashed lines).

To determine which strains from each fecal sample colonized in presence of hCom1, we analyzed the composition of fecal pellets collected in weeks 5-8 to assign species as ‘input’ (hCom1-derived) or ‘invader’ (fecal sample-derived). Using MIDAS, we cannot determine whether a strain present both pre- and post-challenge was derived from hCom1 (i.e., the original strain colonized persistently) or the fecal sample (i.e., a new strain displaced the original strain). To gain further insight into strain displacement versus persistence, we recruited reads from samples taken four weeks post-challenge (week 8) to a database composed of the hCom1 genome sequences, using only reads that were 100% identical to one or more of the genomes. We focused our analysis on genomes with high depth of coverage (≥10X). More than 60% of these strains were covered broadly (≥95%) by perfectly matching reads, indicating that most strains present pre- and post-challenge were either hCom1-derived or a closely related strain (**Figure S2**).

As expected, mice challenged by saline instead of a fecal sample showed no evidence of new species post-challenge (**Figure 2B**). To our surprise, in hCom1-colonized mice challenged by a fecal sample, an average of 89% of the genome copies from week 8 (and 58% of the MIDAS bins, a rough proxy for species) derived from hCom1 (**Figure 2B**). The remaining 11% of the genome copies (and 42% of the MIDAS bins) represent new species that joined hCom1 from one of the fecal samples. Despite the addition of new species, the architecture of the community remained intact (**Figure 2C**): the relative abundances of the hCom1-derived species present post-challenge were highly correlated with their pre-challenge levels (Pearson’s *r* >0.85) (**Figure 2D**). Thus, hCom1 is broadly but not completely resilient to a human fecal challenge.

### Designing and constructing an augmented community

The observation that only a small fraction of the post-challenge community is composed of new strains led us to hypothesize that we could improve the colonization resistance of hCom1 by adding the invading strains, thereby improving its ability to fill niches in the gut. Twenty-five bacterial species entered hCom1 from ≥2 of the 3 fecal samples used as a challenge (**Table S1**); we focused on these species, reasoning that they were more likely to fill conserved niches in the community. We were able to obtain 22/25 from culture collections and we included all of them in the new community. At the same time, we omitted seven species that either failed to colonize initially or were displaced in all three groups of mice (**Figure S3**), reasoning that they were incompatible with the rest of hCom1 or incapable of colonizing the mouse gut under the dietary conditions in which the experiment was performed. Thus, the new community (hCom2) contains 97 strains from hCom1 plus 22 new strains, for a total of 119 (**Figure 3B, Figure S3, Table S2**).

**Figure 3:**
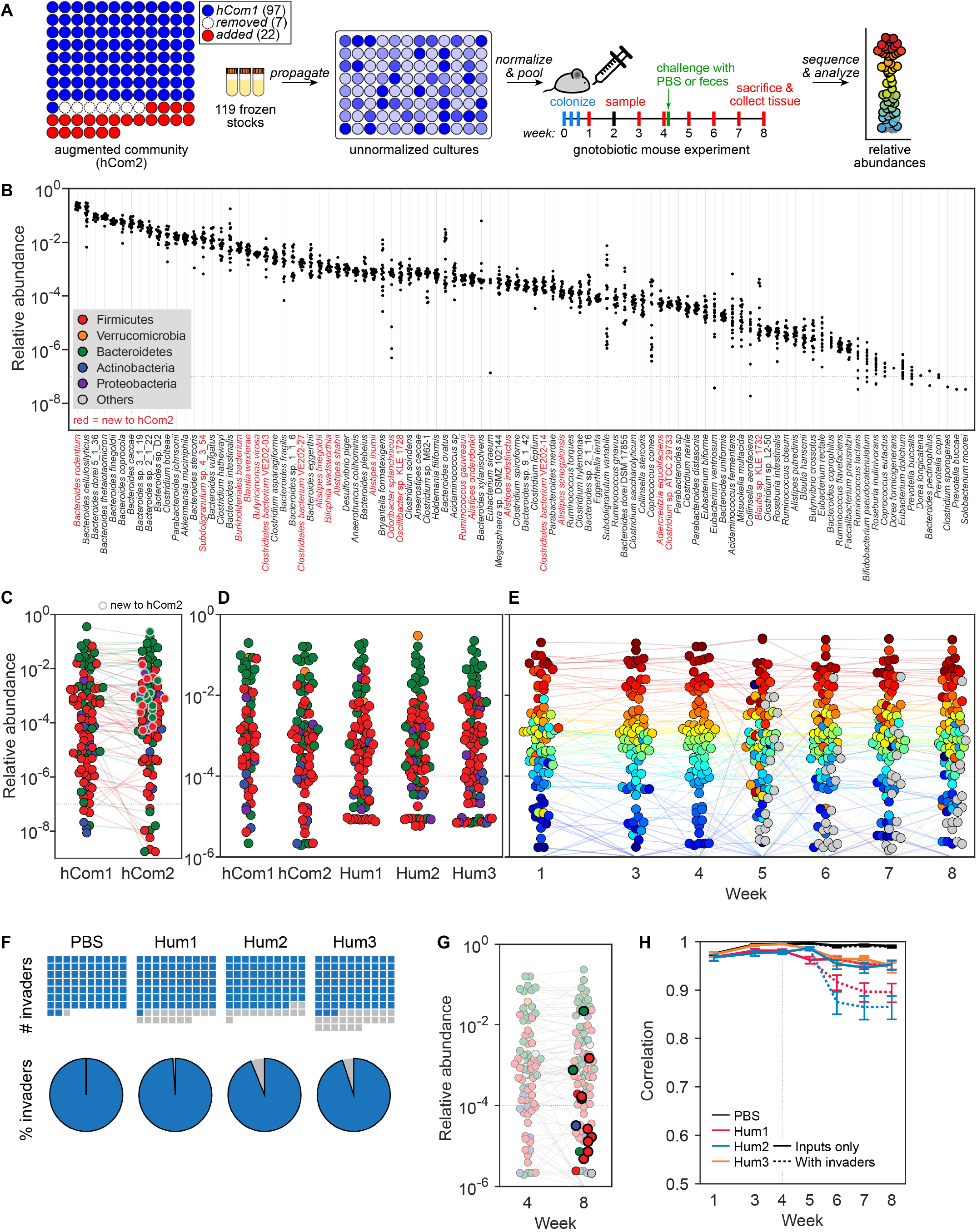
An augmented community with improved resilience to fecal challenge. (**A**) Schematic of the experiment. To augment the community, we added 22 strains that colonized in the presence of hCom1 in ≥2 of the 3 fecal challenge conditions; we also removed seven species that were displaced in all three groups of mice. Thus, the new community (hCom2) contains 97 strains from hCom1 plus 22 new strains for a total of 119. Mice were colonized by hCom2 and housed for four weeks. At the beginning of week 5, they were challenged with the same fecal communities used in the first experiment, or with PBS as a control. After four additional weeks, we used metagenomic sequencing coupled with MIDAS to analyze community composition from fecal pellets collected at weeks 1 and 3-8. (**B**) Comparing the architecture and strain-level relative abundances of hCom1 and hCom2. Each column depicts the relative abundance of an individual strain from hCom2 across all samples at week 4. 100 of the 119 strains were detected; those that are new to hCom2 are colored red. (**C**) Averaged relative abundances of the strains in hCom1 versus hCom2 at week 4. Strains that are new to hCom2 are indicated by a gray outline. Dots are colored by phylum according to the legend in panel B. (**D**) hCom2 resembles a fecal consortium more closely than hCom1. Averaged relative abundances of MIDAS bins—a rough proxy for species—are shown for hCom1- and hCom2-colonized mice versus mice colonized by stool from one of three healthy human donors (Hum1-3). The phylum-level architecture of hCom2 is more closely correlated to that of humanized mice than hCom1 (see **Figure S5**). (**E**) hCom2 is more resilient to fecal challenge than hCom1. Top row: Blue squares in the waffle plots indicate strains that derive from hCom2; gray squares represent strains from the fecal communities. Bottom row: pie charts representing the percentage of MIDAS bins that derive from hCom2 versus the fecal communities. An average of 96% of the genome copies (and 81% of the MIDAS bins) come from hCom2, demonstrating that the resilience of the community was improved markedly by augmentation with strains from the initial challenge. (**F**) The architecture of hCom2 is largely unaffected by fecal challenge. Each dot is an individual strain; the collection of dots in a column represents the community at a single timepoint averaged over the 5 co-housed mice that were challenged with fecal community Hum1. Strains are colored according to their rank-order relative abundance at week 4. Gray circles represent invading species, defined as any species not present in weeks 1-4 in the group of mice shown. (**G**) Nearly all of the invading strains were repeat invaders from the first fecal challenge. The dots representing invading strains are shown in full color; dots representing hCom2-derived strains are partially transparent. Dots that represent repeat invaders from the first fecal challenge experiment have a thick black border. (**H**) The relative abundances of the hCom2-derived species present post-challenge are highly correlated with their pre-challenge levels. Pearson’s correlation coefficient with respect to the average relative abundance in weeks 3 and 4 are shown for the PBS control and 3 fecal community challenges, averaged across mice that received the same challenge. Correlation coefficients are shown for the 119 species in hCom2 (solid lines) and for all species including invaders (dashed lines).

We constructed hCom2 by culturing each strain separately and then mixing cultures after normalization. We colonized four groups of germ-free SW mice with hCom2, collecting fecal pellets weekly. As before, we measured community composition by metagenomic sequencing with NinjaMap (**Figure 3A**). The gut communities of hCom2-colonized mice rapidly reached a stable configuration (Pearson’s *r* with respect to week 8 >0.97) (**Figure S4**). 100 of the 119 strains were above the limit of detection; hCom1-derived strains colonized at similar relative abundances in the context of the augmented community (with similarly low CVs across mice). The strains that were new to hCom2 exhibited a wide range of relative abundances; *Bacteroides rodentium* became the most abundant species, whereas the least abundant of the new species, *Blautia sp*. KLE 1732, had a mean abundance ∼10^−4^ (**Figure 3B**). Moreover, the architecture of the community more closely resembled that of a human fecal consortium at both the phylum and strain level (**Figures 3D, S5**).

### The augmented community is more resilient to human fecal challenge

Our goal in constructing hCom2 was to improve its completeness as assessed by its ability to occupy niches in the gut. To test whether hCom2 is more complete than hCom1, we challenged hCom2-colonized mice at the beginning of week 5 with the same fecal samples used to challenge hCom1, enabling us to compare results between the challenge experiments. Importantly, the 22 strains used to augment hCom1 were obtained from culture collections rather than the fecal samples themselves, reducing the likelihood that hCom2 and the fecal samples have overlapping membership at the strain level (Garud et al. 2019). Indeed, by recruiting sequencing reads to the genomes of the new organisms in hCom2, we found that 17/22 were covered broadly (≥95%) by perfectly matching reads, consistent with the view that they were derived from hCom2 and not the fecal challenge (**Figure S6**).

An average of 96% of the genome copies (and 81% of the MIDAS bins) from week 8 derived from the strains in hCom2, demonstrating that the colonization resistance of hCom2 is markedly improved over hCom1 (**Figure 3E**). The remaining 4% of reads (and 19% of MIDAS bins) represent species that engrafted in spite of the presence of hCom2 (**Figures 3E, S7**). Strikingly, nearly all of the species that invaded hCom2 also invaded hCom1; we were either unable to obtain an isolate for inclusion in hCom2 or the species invaded hCom1 from only 1 of the 3 fecal samples used as a challenge, falling below our threshold for inclusion. These species represented virtually all of the remaining genome copies. We conclude that more extensive augmentation—based on the results of the first challenge experiment—would likely have enhanced colonization resistance beyond 99%.

Moreover, compared to hCom1, the composition of hCom2 post-challenge was more similar to its pre-challenge state (Pearson’s *r* relative to week 4 >0.95, **Figure 3F-G**). Taken together, these data show that hCom2 is more stable and complete than hCom1, and that the backfill process is robust and fault-tolerant in identifying species that can occupy unfilled niches.

### Reproducibility of colonization

Our original goal in building a complex defined community was to develop a model system for the gut microbiome. Having demonstrated that hCom2 is stable and resilient to invasion, we sought to assess whether it has the attributes of a model system.

A threshold requirement is a substantial degree of reproducibility. We started by analyzing data from the second fecal challenge experiment in order to assess the technical reproducibility of community composition in mice colonized by hCom2. At week 4, strain abundances in 20 mice across 4 cages colonized by the same hCom2 inoculum were highly similar (pairwise Pearson’s correlation coefficients 0.96±0.01, **Figure S8**).

Biological reproducibility was a greater concern. Given the complexity of hCom1 and hCom2, variability in the growth of individual strains could lead to substantial differences in the composition of inocula constructed on different days. To determine the extent to which this variability affects community architecture *in vivo*, we compared community composition in four groups of mice colonized by replicates of hCom2 constructed independently on different days (**Figure 4A-B**). The communities displayed a striking degree of similarity in relative abundance profiles after 4 weeks (Pearson’s correlation coefficient >0.95 between all pairs of biological replicates). We conclude that a relatively constant nutrient environment enables input communities with widely varying relative abundances to reach the same steady state configuration, consistent with ecological observations in other microbial communities (Hibberd et al. 2017; Goldford et al. 2018; Aranda-Díaz et al. 2020; Venturelli et al. 2018). This high degree of biological reproducibility will be enabling for the use of complex defined communities as experimental models.

**Figure 4:**
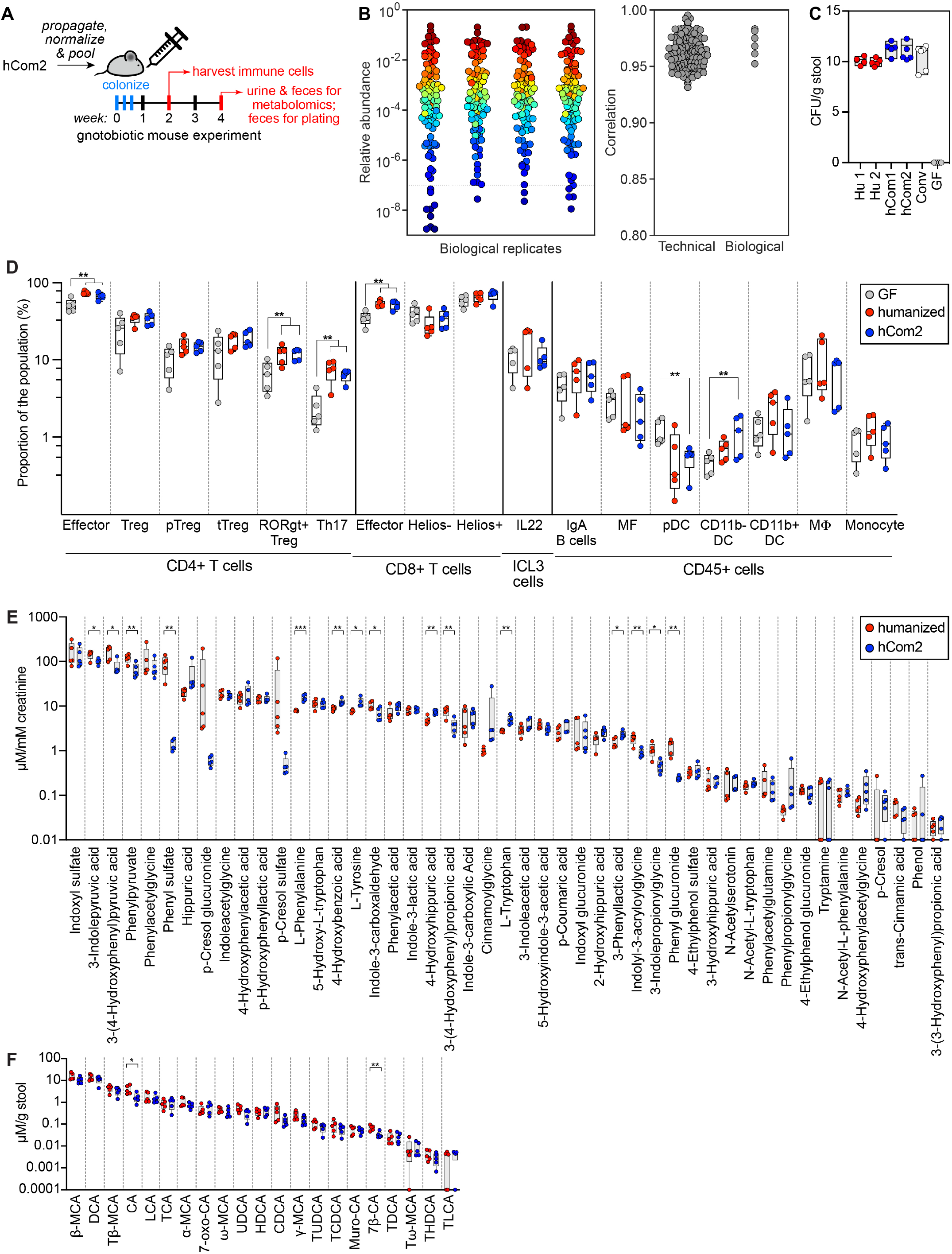
hCom2-colonized mice are phenotypically similar to humanized mice. (**A**) Schematic of the experiment. Germ-free SW mice were colonized with hCom2 or a fecal sample from a healthy human donor. One cohort of mice was sacrificed at two weeks for immune cell profiling; another was sacrificed at four weeks for targeted metabolite analysis. (**B**) The architecture of hCom2 in mice is highly reproducible. Left: community composition is highly similar across four biological replicates. Each dot is an individual strain; the collection of dots in a column represents the community at 4 weeks averaged over 5 mice co-housed in a cage. Strains are colored according to their average rank-order relative abundance across all samples. Right: Pearson’s pairwise correlation coefficients for technical and biological replicates. (**C**) hCom2, hCom1, and humanized mice have similar bacterial cell densities *in vivo*. Fecal samples from hCom2-colonized, hCom1-colonized, humanized, specific pathogen-free (SPF), or germ-free (GF) mice were homogenized and plated anaerobically on Columbia Blood Agar to enumerate colony forming units. (**D**) Immune cell types and numbers were broadly similar between hCom2-colonized and humanized mice and distinct from germ-free mice. Colonic immune cells were extracted from hCom2-colonized, humanized, or germ-free mice (all C57BL/6), stained for cell surface markers, and assessed by flow cytometry. Statistical significance was assessed using a Student’s two tailed t-test (**: *p*<0.05). (**E**) hCom2-colonized mice and humanized mice have a similar profile of microbiome-derived metabolites. Urine samples from hCom2-colonized and humanized mice were analyzed by targeted metabolomics to measure a panel of aromatic amino acid metabolites by LC-MS. Statistical significance was assessed using a Student’s two tailed t-test (*: *p*<0.05; **: *p*<0.001). (**F**) Bile acids were extracted from fecal pellets collected from hCom2-colonized and humanized mice and were quantified by LC-MS. Statistical significance was assessed using a Student’s two tailed t-test (*: *p*<0.05; **: *p*<0.001).

To further investigate the potential for hCom2 to function as a model microbiome, we assessed its composition in a second strain of mice. Since the experiments to develop hCom2 used outbred SW mice, we chose 129/SvEv, an inbred strain derived from the C57BL/6 background. We colonized germ-free 129/SvEv mice with hCom2 and collected fecal pellets after 4 weeks of colonization. Community composition was highly correlated with that of SW mice (Pearson correlation coefficient >0.95) (**Figure S8B**). These data indicate that hCom2, like the human gut microbiome (Rothschild et al. 2018), is robust to changes in host genotype.

### hCom2-colonized mice are phenotypically similar to humanized mice

We performed three additional experiments to determine the degree to which hCom2-colonized mice resemble humanized mice. Since our defined communities are composed of human fecal isolates, we colonized germ-free mice with hCom2 or an undefined human fecal community and assayed phenotypes after 4 weeks (**Figure 4A**). First, fecal pellets from each mouse were serially diluted and plated on Columbia blood agar to estimate the bacterial cell density in each community. Each group contained 10^11^-10^12^ colony forming units per gram of feces (**Figure 4C**), similar to previously reported estimates from humans and from conventional and humanized mice (Vandeputte et al. 2017; Ley et al. 2006). Thus, hCom2 colonizes the mouse gut to a similar extent as a normal murine or human fecal community.

Next, we sought to determine whether mice colonized by hCom2 harbor a similar immune cell profile to that of humanized mice. We extracted and stained colonic immune cells and assayed them by flow cytometry. Most immune cell subtypes were similarly abundant in humanized and hCom2-colonized mice (**Figure 4D**), indicating that—at least in broad terms—hCom2-colonized mice are immunologically comparable to humanized mice.

Finally, to determine whether hCom2-colonized mice had a similar profile of microbiome-derived metabolites to humanized mice, we analyzed fecal pellets and urine samples using targeted metabolomics. Aromatic amino acid metabolite levels in urine (**Figure 4E**) and primary and secondary bile acid levels in feces (**Figure 4F**) were comparable between hCom2-colonized and humanized mice. Notably, hCom1-colonized mice exhibited lower levels of secondary bile acids than hCom2-colonized mice, indicating that some of the species used to augment hCom2 likely contribute to bile acid metabolism. Taken together, these data suggest that hCom2 is a reasonable model of gut microbial metabolism. A more thorough study would be needed to reveal finer-scale differences among mice humanized by distinct fecal samples.

### hCom2 exhibits robust colonization resistance against pathogenic *Escherichia coli*

To demonstrate the utility of hCom2 as a model system, we used it to study an emergent property of gut communities: their ability to resist colonization by pathogens and pathobionts (Buffie et al. 2015). To test whether hCom2 has this property, we studied invasion by enterohemorrhagic *Escherichia coli* (EHEC). We chose this strain for three reasons. First, EHEC is responsible for life-threatening diarrheal infections and hemolytic uremic syndrome, and enteric colonization by other *E. coli* strains has been linked to malnutrition and inflammatory bowel disease (Palmela et al. 2018; Pham et al. 2019). Second, colonization resistance to *E. coli* and other Enterobacteriaceae has been studied in detail (Stromberg et al. 2018; Velazquez et al. 2019; Litvak et al. 2019), but the commensal strains responsible and mechanisms by which they act are incompletely understood. Finally, hCom2 harbors no Enterobacteriaceae and only two other (distantly related) strains of Proteobacteria, *Desulfovibrio piger* and *Bilophila wadsworthia*, so resistance to *E. coli* colonization would require a mechanism other than exclusion by a close relative occupying the same niche.

To test whether hCom2 is capable of resisting EHEC engraftment, we colonized germ-free SW mice with hCom2 or one of two other communities: a 12-member community (12Com) similar to one used in previous studies (McNulty et al. 2013) or an undefined stool sample from a healthy human donor (**Figure 5A**). hCom2 and 12Com do not contain any Enterobacteriaceae; to test whether non-pathogenic Enterobacteriaceae enhance colonization resistance to EHEC, we colonized two additional groups of mice with variants of hCom2 and 12Com to which a mixture of seven non-pathogenic Enterobacteriaceae strains were added (six *E. coli* and *Enterobacter cloacae*). After four weeks, we challenged with EHEC and assessed invasion by selective plating under aerobic growth conditions (**Figure 5A**).

**Figure 5:**
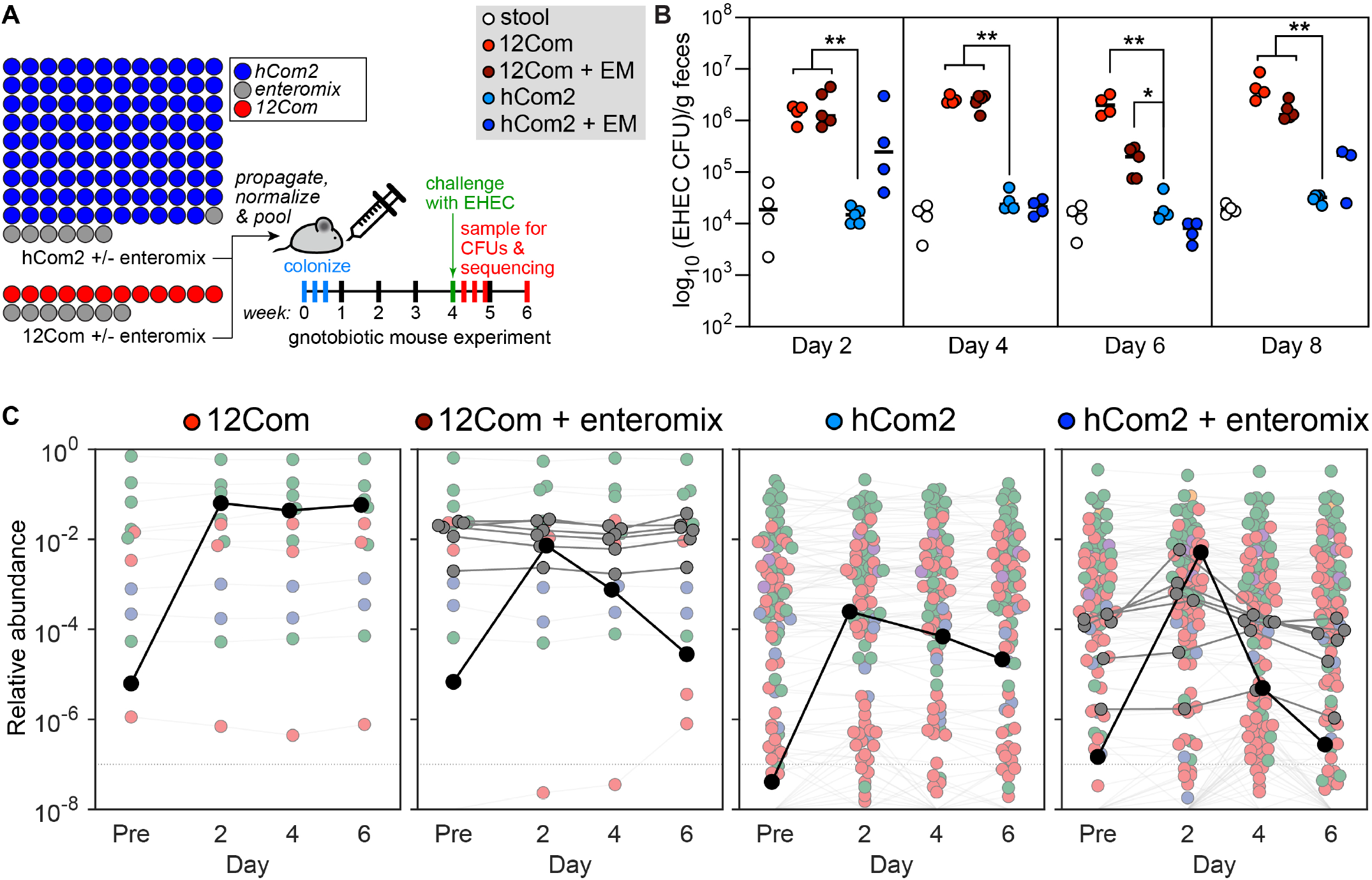
hCom2 exhibits colonization resistance against enterohemorrhagic *E. coli*. (**A**) Schematic of the experiment. We colonized germ-free SW mice with hCom2 or one of two other communities: a 12-member synthetic community (12Com) or a fecal community from a healthy human donor. hCom2 and 12Com do not contain any Enterobacteriaceae; to test whether non-pathogenic Enterobacteriaceae enhance colonization resistance to EHEC, we colonized two additional groups of mice with variants of hCom2 and 12Com to which a mixture of seven non-pathogenic Enterobacteriaceae strains were added (six *E. coli* and *Enterobacter cloacae*, enteromix (EM)). After four weeks, we challenged with 10^9^ colony forming units of EHEC and assessed the degree to which it colonized in two ways: by EHEC-selective plating under aerobic growth conditions, and by metagenomic sequencing with NinjaMap analysis. (**B**) hCom2 exhibits a similar degree of EHEC resistance to that of a fecal community in mice. Colony forming units of EHEC in mice colonized by the four different communities are shown. As expected, the fecal community conferred robust colonization resistance while 12Com did not. Despite lacking Enterobacteriaceae, hCom2 exhibited a similar level of EHEC resistance to that of an undefined fecal community; the addition of non-pathogenic Enterobacteriaceae further improved this phenotype. (**C**) The architecture of hCom2 is stable following EHEC challenge. Each dot is an individual strain; the collection of dots in a column represents the community at a single timepoint averaged over four co-housed mice. Strains are colored according to their phylum; EHEC is shown in black and members of the enteromix community are shown in gray.

Consistent with previous reports (Mohawk and O’Brien 2011; Stromberg et al. 2018), the undefined human fecal sample conferred robust resistance against EHEC colonization (**Figure 5A**). In contrast, 12Com allowed much higher levels of EHEC growth; the addition of non-pathogenic Enterbacteriaceae improved the phenotype but did not restore full EHEC resistance (**Figure 5A, B**). Despite lacking Enterobacteriaceae, hCom2 exhibited a similar level of EHEC resistance to that of an undefined fecal community; the addition of non-pathogenic Enterobacteriaceae further improved this phenotype (**Figure 5A**). Thus, hCom2 is sufficiently complete to exhibit comparable levels of colonization resistance to a native fecal community.

## DISCUSSION

The process by which we augmented a defined community revealed two unexpected findings. First, a community composed of strains from >100 distinct donors can be stable *in vivo*. It remains to be seen whether there are appreciable differences in stability—or in fine-scale genomic and phenotypic adaptation—between communities composed of isolates from a single donor (in which strains have co-existed for years) versus multiple donors (in which strains have no prior history together).

Second, the process we introduce here for filling open niches is surprisingly robust and fault tolerant. Most notably, nearly all of the strains that invaded hCom2 upon fecal challenge had previously invaded hCom1, indicating that niche filling is deterministic. Importantly, the backfill process caused little perturbation to the structure of the existing community, suggesting that it will result in a progressive improvement of the community. While the backfill process can only fill niches that are conserved from mice to humans, the observation that most of our human strains engrafted suggests that many niches are conserved.

If we broaden our strain inclusion criteria, there is a reasonable likelihood we could have achieved >99% colonization resistance after just one round of augmentation. To further enhance niche filling and stability, it would help to subject hCom2 to further rounds of backfill involving fecal samples from other donors, ideally in the presence of a varying diet. It might also be possible to improve niche occupancy, e.g., in the setting of intestinal inflammation by performing the backfill process in a murine model of inflammatory bowel disease.

There is a great need for a common model system for the gut microbiome that is completely defined and complex enough to capture much of the biology of a full-scale community. We show that hCom2 is a reasonable starting point for such a system: in spite of its complexity, it colonizes mice in a highly reproducible manner. Moreover, hCom2 faithfully models the carrying capacity, immune cell profile, and metabolic phenotypes of humanized mice. There remain some modest differences in metabolic and immune profiles, and the community is still missing certain taxa that will likely be important to add. Nonetheless, taken together, our findings suggest that hCom2 is a reasonable starting point for a model system of the gut microbiome.

One of the most interesting possibilities for such a system would be to enable reductionist experiments downstream of a community transplantation experiment (e.g., to identify strains responsible for a microbiome-linked phenotype). Although we did not identify the strains responsible for colonization resistance to EHEC, follow-up experiments in which one or several strains at a time are eliminated from the community could narrow this down from the phylum level to individual strains. Efforts to identify the strains responsible for other microbiome-linked phenotypes including response to cancer immunotherapy, caloric harvest, and neural development, would be of great interest.

Our study has two important limitations. Since we challenged hCom2 with the same fecal samples used to challenge hCom1, we do not yet know whether hCom2 is universally stable to challenge by any fecal sample. However, in light of the fact that fecal communities from healthy adults are typically functionally replete and ecologically stable, our data demonstrating resilience to challenge by three such samples is a foundational starting point for the development of a defined community that is stable to broad set of microbial and environmental perturbations.

Second, it is unclear how many more bacterial strains (or other components) may be necessary to model the full functional capacity of a native human microbiome. Prior estimates of the number of species in a typical human microbiome range from ∼150-300 (Faith et al. 2013; Kraal et al. 2014; Qin et al. 2010); in addition, hCom2 does not contain any archaea, fungi, or viruses, which could be added to broaden its diversity. Nonetheless, the observation that a defined community of just 119 strains exhibits remarkable stability bodes well for future efforts. We estimate that hCom2 is within 2-fold of native-scale complexity, so a full-scale system is experimentally feasible. As a starting point for efforts to build such a system, hCom2 will provide a standard for assessing the genomic and functional completeness of model communities, with the ultimate goal of modeling native-scale human microbiomes.

### STAR*METHODS

Detailed methods are provided in the online version of this paper and include the following:

- KEY RESOURCES TABLE
- RESOURCE AVAILABILITY
  - Lead contact
  - Materials availability
  - Data and code availability
- EXPERIMENTAL MODEL AND SUBJECT DETAILS
  - Bacterial strains and culture conditions
  - Preparation of 12Com
  - Preparation of Enteromix
  - Collection and preservation of human stool
  - Preparation of human stool
- METHOD DETAILS
  - Metagenomic sequencing
  - Augmenting the NinjaMap database
  - Metagenomic read mapping
  - Backfill experiment
  - Reproducibility and colonization experiments
  - Bacterial load estimates
  - Immune profiling
  - Metabolomics
  - *E. coli* colonization resistance
- QUANTIFICATION AND STATISTICAL ANALYSIS

#### Lead contact

Further information and requests for resources and reagents should be directed to and will be fulfilled by the lead contact, Michael Fischbach.

#### Materials availability

*C. sporogenes* strains are available on request. The strains used in this study are available from the sources listed in the Key Resource Table.

#### Data and code availability

Original metagenomic and whole-genome sequencing data generated for this study will be available at the time of publication. The analysis tool NinjaMap is available at https://github.com/czbiohub/NinjaMap.

### EXPERIMENTAL MODEL AND SUBJECT DETAILS

#### Bacterial strains and culture conditions

Bacterial strains were selected based on HMP sequencing data (Kraal et al. 2014). We obtained all species from publicly available repositories; the origin of each strain is listed in the Key Resources Table. Strains were cultured anaerobically in a 10% CO_2_, 5% H_2_, and 85% N_2_ atmosphere in autoclave-sterilized 2.2-mL 96-well deep well plates (Thomas Scientific, Cat. #1159Q92). To create frozen stocks in 96-well format, strains were aliquoted 1:1 into sterile 50% glycerol, capped with a silicone fitted plate mat (Thomas Scientific, Cat. #SMX-DW96S20), and the edges were sealed with oxygen-impervious yellow vinyl tape (Coy Labs, Cat. #1600330w).

#### Preparation of synthetic community

For all germ-free mouse experiments, strains were cultured and pooled in the following manner: From frozen stocks in 96-well plates, 100 µL of each strain was used to inoculate 900 µL of autoclave-sterilized media of the appropriate type for each strain in 2.2-mL 96-well deep well plates. Strains were diluted 1:10 every 24 h for 2 days into fresh growth medium in 2.2-mL deep well plates, and then diluted 1:10 into 4 mL of the appropriate medium in 5-mL 48-well deep well plates (Thomas Scientific, Cat. #1223T83). After 24 h, the optical density at 600 nm (OD_600_) of each well was measured. Based on measurements of absorbance at 600 nm and enumeration of colony forming units (CFUs), we found that an OD_600_ of 1.3 corresponds to ∼10^9^ cells/mL for *E. coli*. Using this estimate, we pooled appropriate volumes of each culture corresponding to 2 mL at OD_600_=1.3, centrifuged for 5 min at 5000 x *g*, and resuspended the pellet into 2 mL of 20% glycerol that had been pre-reduced for at least 48 h. For each inoculum preparation cycle, a small number of strains (<20) typically did not reach OD_600_∼1.3. For these strains, the entire 4 mL culture volume was used for pooling. Volumes were scaled up accordingly if more inoculum was required for an experiment. Following pooling and preparation, 1.2 mL of the synthetic community was aliquoted into 2 mL Corning cryovials (Corning, Cat. #430659), removed from the anaerobic chamber, and transported to the vivarium where each vial was uncapped and its contents orally gavaged into mice within 1 min of uncapping. Each mouse received 200 µL of the mixed community inoculum. For the initial backfill experiments, we used freshly prepared inoculum; for all subsequent experiments, the inoculum was frozen in cryovials at -80 °C; on the day of the experiment, it was defrosted and administered by oral gavage. The target for the inoculation procedure was that each mouse receive ∼10^8^ cells of each bacterial strain in a 200 µL volume, for a total of ∼10^10^ bacterial cells since hCom1 and hCom2 harbor 104 and 119 strains, respectively.

#### Preparation of 12Com

Cultures of the 12 strains in 12Com (*Bacteroides thetaiotaomicron* VPI-5482, *Bacteroides caccae* ATCC 43185, *Bacteroides ovatus* ATCC 8483, *Bacteroides uniformis* ATCC 8492, *Bacteroides vulgatus* ATCC 8482, *Clostridium scindens* ATCC 35704, *Collinsella aerofaciens* ATCC 25986, *Dorea longicatena* DSM 13814, *Eggerthella lenta* DSM 2243, *Eubacterium rectale* ATCC 33656, *Parabacteroides distasonis* ATCC 8503, and *Ruminococcus torques* ATCC 27756) were prepared in their respective growth media and propagated anaerobically for 24 h to OD_600_∼1.3. 2 mL of each strain was pooled, centrifuged for 5 min at 5000 x *g*, and the pellet was resuspended in 2 mL of 20% pre-reduced glycerol and frozen in 1 mL aliquots in 2 mL Corning cryovials.

#### Preparation of Enteromix

Six strains of non-pathogenic *Escherichia coli* (strains MITI 27, MITI 117, MITI 135, MITI 139, MITI 255, MITI 284) and one strain of *Enterobacter cloacae* (MITI 173) were isolated from the stool of a healthy human donor by mass spectrometry-guided enrichment culture (Lagier et al. 2016). Strains were stored at -80 °C in 25% glycerol. To prepare cultures for mouse colonization, strains were grown overnight in BHI broth, diluted 1:10 into 5 mL BHI broth, and cultured to OD_600_ ∼1.3. 2 mL of each strain were pooled, centrifuged for 5 min at 5000 x *g*, and the pellet was resuspended in 200 µL of 20% pre-reduced glycerol. 100 µL of this mixture were added to a tube containing 1 mL of previously prepared hCom2 or 12Com inoculum to create hCom2+Enteromix or 12Com+Enteromix, respectively. Each mouse was orally gavaged with 220 µL of the appropriate community. The estimated amount of each Enteromix strain administered to mice was 10^9^ cells per 20 µL dose.

#### Collection and preservation of human stool

For all experiments, human stool was preserved in the same manner for inoculation into germ-free or hCom1/2-colonized mice. Specifically, freshly voided human stool was collected in a sterile container and transported into the anaerobic chamber within 5-10 min. The stool was weighed, mixed 1:1 with an equivalent volume of pre-reduced PBS, and stored at -80 °C.

#### Preparation of human stool

For human fecal challenge experiments, a stool mixture was defrosted in the anaerobic chamber and diluted 1:100 into pre-reduced PBS. 1 mL was aliquoted into pre-reduced 2 mL Corning cryovials, removed from the anaerobic chamber, and transported to the vivarium, where each vial was uncapped and orally gavaged into mice within 1 min of uncapping. Each mouse received 200 µL of the bacterial mixture. Stool contains ∼10^11^ colony forming units per gram of feces (Vandeputte et al. 2017); based on the dilutions performed, we estimate that each mouse received 10^8^-10^10^ bacterial cells in the fecal challenge.

For all non-challenge stool colonization experiments, the preserved stool mixture was defrosted in the anaerobic chamber and diluted 1:2 into pre-reduced PBS. 1 mL of the resulting mixture was aliquoted into pre-reduced 2 mL Corning cryovials, removed from the anaerobic chamber, and transported to the vivarium, where each vial was uncapped and orally gavaged into mice within 1 min of uncapping. Each mouse received 200 µL of the bacterial mixture, equivalent to 10^10^-10^11^ bacterial cells per mouse.

#### Metagenomic sequencing

The same experimental pipeline was used for sequencing bacterial isolates and microbial communities. Bacterial cells were pelleted by centrifugation in an anaerobic environment. Genomic DNA was extracted using the DNeasy PowerSoil HTP kit (Qiagen) and the quantity of extracted genomic DNA was measured in a 384-well format using the Quant-iT PicoGreen dsDNA Assay Kit (Thermo Fisher). Sequencing libraries were generated in a 384-well format using a custom, low-volume protocol based on the Nextera XT process (Illumina). Briefly, the DNA concentration from each sample was normalized to 0.18 ng/µL using a Mantis liquid handler (Formulatrix). In cases where concentration was below 0.18 ng/µL, the sample was not diluted further. Tagmentation, neutralization, and PCR steps of the Nextera XT process were performed on the Mosquito HTS liquid handler (TTP Labtech), creating a final volume of 4 µL per library. During the PCR amplification step, custom 12-bp dual unique indices were introduced to eliminate barcode switching, a phenomenon that occurs on Illumina sequencing platforms with patterned flow cells (Costello et al. 2018). Libraries were pooled at the desired relative molar ratios and cleaned up using Ampure XP beads (Beckman) to effect buffer removal and library size selection. The cleanup process was used to remove fragments shorter than 300 bp and longer than 1.5 kbp. Final library pools were quality checked for size distribution and concentration using the Fragment Analyzer (Agilent) and qPCR (BioRad). Sequencing reads were generated using the NovaSeq S4 flow cell or the NextSeq High Output kit, both in 2×150 bp configuration. 5-10 million paired-end reads were targeted for bacterial isolates and 20–30 million paired end reads for bacterial communities.

#### Augmenting the NinjaMap database

We obtained the latest available RefSeq (O’Leary et al. 2016) assembly for each strain in our communities and assessed their quality based on contig statistics from Quast v. 5.0.2 (Gurevich et al. 2013) and SeqKit v. 0.12.0 (Shen et al., 2016), and GTDB-tk v. 1.2.0 (Chaumeil et al. 2019) for taxonomic classification. A ‘combination score’ was calculated as a linear combination of the completeness and contamination scores (completeness–5×contamination) derived from the CheckM v. 1.1.2 lineage workflow (Parks et al. 2015); such a score has been used previously, along with the metrics described here (https://gtdb.ecogenomic.org/faq#gtdb_selection_criteria), to include/exclude genomes in the GTDB release 89 database (Parks et al. 2020; Parks et al. 2018). Genomes that contained any number of Ns, contained over 100 contigs, contained GTDB lineage warnings or multiple matches, or had CheckM completeness <90, contamination >10, and combination score <90 were selected for resequencing and reassembly.

Our hybrid assembly pipeline contains a workflow for *de novo* or reference-guided genome assembly using Illumina short reads in combination with long reads from PacBio or Nanopore. The workflow has three main steps: read pre-processing, hybrid assembly, and contig post-processing. Read pre-processing included 1) quality trimming/filtering (bbduk.sh adapterFile=“adapters,phix” k=23, hdist=1, qtrim=rl, ktrim=r, entropy=0.5, entropywindow=50, entropyk=5, trimq=25, minlen=50), with adapters and phix removed with kmer right-trimming, kmer size of 23, Hamming distance 1 (allowing one mismatch), quality trimming of both sides of the read, filtering of reads with average entropy <0.5 with entropy kmer length of 5 and a sliding window of 50, trimming to a Q25 quality score, and removal of reads with length <50 bp; 2) deduplication (bbdupe.sh); 3) coverage normalization (bbnorm.sh min=3) such that depth <3x was discarded; 4) error correction (tadpole.sh mode=correct); and 5) sampling (reformt.sh). All pre-processing was carried out using BBtools v. 38.37 for short reads. For long reads, we used filtlong v. 0.2.0 (fitlong --min_length 1000 --keep_percent 90 --length_weight 10) to discard any read <1 kb and the worst 10% of read bases, as well as to weight read length as more important when choosing the best reads. Hybrid assembly was performed by Unicycler v. 0.4.8 (Wick et al. 2017) with default parameters using pre-processed reads. After assembly, the contigs from the assembler were scaffolded by LRScaf v. 1.1.9 (Qin et al. 2018) with default parameters. If the initial assembly did not produce the complete genome, gaps were filled using long reads by TGS-GapCloser v. 1.0.1 (Xu et al. 2019) with default parameters.

If no long reads were available, short paired-end reads were *de novo*-assembled using SPAdes v. 3.13.1 (Bankevich et al. 2012) with the --careful option to reduce the number of mismatches and short indels during assembly of small genomes. Assembly quality was assessed based on the CheckM v. 1.1.2 lineage. If contamination was detected, the assembly was binned from the contaminated assembly using MetaBAT2 v. 2.2.14 (Kang et al. 2019) with default parameters.

Finally, assembled genomes were evaluated using the same criteria as the RefSeq assemblies, and the assembly for each species with better overall quality metrics was chosen as the reference assembly. This procedure resulted in the replacement of 85 genomes: two obtained from a PacBio/Illumina hybrid assembly, 69 from a Nanopore/Illumina hybrid assembly, one from a reference-guided Illumina assembly, and seven from short-read assemblies of the respective isolate samples followed by binning (**Table S3**).

#### Metagenomic read mapping

Paired-end reads from each sample were aligned to the hCom1 or hCom2 database using Bowtie2 with maximum insert length (-maxins) set to 3000, maximum alignments (-k) set to 300, suppressed unpaired alignments (--no-mixed), suppressed discordant alignments (--no-discordant), suppressed output for unaligned reads (--no-unal), required global alignment (--end-to-end), and using the “--very-sensitive” alignment preset (command: --very-sensitive -maxinsX 3000 -k 300 --no-mixed --no-discordant --end-to-end --no-unal). The output was piped into Samtools v. 1.9 (Li et al. 2009), which was used to convert the alignment output from SAM output stream to BAM format and then sort and index the BAM file by coordinates. Alignments were filtered to only keep those with >99% identity for the entire length of the read.

The median percentage of unaligned reads was 4.95% (range 4.10% - 8.35%). To assess the origin of these reads, we performed a BLAST v2.11.0+ search through the ncbi/blast:latest docker image with parameters “-outfmt ‘6 std qlen slen qcovs sscinames staxids’ -dbsize 1000000, -num_alignments 100” from a representative sample against the ‘NCBI - nt’ database as on 2021-02-16. We then filtered the BLAST results to obtain the top hits for a given query. Briefly, the script defined top hits as ones that had an e-value <= 1e-30, percent identity >= 99% and were within 10 percent of the best bit score for that query. To visualize and summarize the output, we used the ktImportTaxonomy script from the Krona package with default parameters. Reads were aggregated by NCBI taxon id and separately by genus. We found that most of the hits are from taxa that are closely related to the organisms in our community, while others are from the mouse genome. We conclude that our experiments did not suffer from any appreciable level of contamination.

#### Backfill experiment

Individual strains were cultured in their respective media (**Table S2**), normalized, and pooled to form the synthetic community as described in ‘Preparation of bacterial synthetic community.’ Mice were orally gavaged with freshly prepared synthetic community three days in a row and were sampled weekly for 4 weeks. After 4 weeks, mice were orally gavaged with stool from one of three healthy human donors (one donor per 5 mice) or PBS as a control.

#### Reproducibility and colonization experiments

Groups of five 6-8-week-old female germ-free SW mice were colonized for 4 weeks with hCom1 or hCom2 and fecal pellets were sampled after 4 weeks. These fecal pellets were subjected to DNA extraction, metagenomic sequencing, and NinjaMap read mapping to estimate strain relative abundances.

#### MIDAS analyses

MIDAS was run using the database v. 1.2 with default parameters on each library. Results were aggregated and filtered to include only MIDAS species representatives (buckets) that were called based on ≥2 reads and reported a minimum of 10^−4^ relative abundance. To determine which invading species to use in augmenting hCom1, a threshold relative abundance of 10^−4^ was applied. A species was selected to augment hCom1 if it is present above the threshold in 2-3 of the 3 challenge groups.

#### MIDAS sensitivity analysis

To determine the sensitivity of MIDAS for analyses of strains in our communities, we generated error-free 150-bp paired-end reads *in silico* for each genome. Each simulated read set was individually processed by MIDAS. While most genomes were identified correctly and assigned to a single MIDAS bucket, 22 strains from hCom1 and hCom2 cross-mapped to multiple buckets. As expected, MIDAS was unable to separate closely related strains, with 14 MIDAS buckets from hCom1 and 17 from hCom2 recruiting reads from more than one strain (**Table S4**).

#### Analyzing strain displacement versus persistence

To determine the coverage of genomes from hCom1 and hCom2 in week 8 samples after a fecal challenge, reads were aligned to two Bowtie2 databases, hCom1 (version SCv1.2) and hCom2 (version SCv2.3). Each alignment file was filtered to only include alignments with 99% or 100% identity at 100% alignment length. Alignments at 99% identity were performed to recruit reads from any strain that was very similar but not identical. The breadth of coverage—i.e., the percentage of the genome covered by at least 1 read—and the depth of coverage (the average number of reads covering positions in the genome) was calculated for each organism in each sample at both identity thresholds.

Results from the MIDAS analysis of each sample were combined with MIDAS bucket strain contributions from the sensitivity analysis and strain coverage metrics. Most of the high abundance strains had high coverage depth and breadth of coverage at 99% and 100% identity, suggesting that the original strains (or highly similar variants) were present in the samples at week 8.

#### Bacterial load estimates

6-8-week-old female germ-free SW mice were colonized for 4 weeks with hCom1, hCom2, or one of two human stool samples, and fecal pellets were sampled after 4 weeks. Female germ-free and conventional SW mice of the same age were sampled at the same time. Each colonization cohort contained 5 mice. For each mouse, two fecal pellets were collected in a pre-weighed 1.5 mL Eppendorf tube containing 200 µL of transport medium. After collection and weighing, the mass of the tube prior to sampling was subtracted to calculate stool weight. Samples were transferred into the anaerobic chamber and each pellet was crushed with a 1 mL pipette tip and vortexed at maximum speed for 30 s to create a homogenous mixture. This mixture was serially diluted 1:10 twelve successive times; each dilution was plated on pre-reduced Columbia blood agar plates and incubated at 37 °C. After 24 h, colonies were counted for each dilution. Fecal pellets were also subjected to DNA extraction, metagenomic sequencing, and NinjaMap analysis to estimate strain relative abundances.

#### Immune profiling

6-8-week-old female germ-free C57BL/6 mice were colonized for 2 weeks with hCom2, a human stool sample, or PBS as a negative control and fecal pellets were collected after 2 weeks. Mice were then sacrificed, colonic tissue was dissected, and immune cells were isolated using the Miltenyi Lamina Propria kit and Gentle MACS dissociator. Immune cells were stained using the antibodies listed in the Key Resource Table at 1:200 dilution and assessed using a LSRII flow cytometer. Fecal pellets were subjected to DNA extraction, metagenomic sequencing, and NinjaMap analysis to estimate strain relative abundances.

#### Metabolomics

Cohorts of 6-8-week-old female germ-free SW mice were colonized for 4 weeks with hCom1, hCom2, or one of two human stool samples; urine and fecal pellets were sampled after 4 weeks. Female germ-free and conventional SW mice of the same age were sampled at the same time. Fecal pellets were subjected to DNA extraction, metagenomic sequencing, and NinjaMap analysis to estimate strain relative abundances.

#### Sample preparation for LC/MS analysis

For urine samples, 5 µL of urine was diluted 1:10 with ddH_2_O and mixed with 50 µL of internal standard water solution (20 μM 4-chloro-L-phenylalanine and 2 μM d^4^-cholic acid). After centrifugation for 15 min at 4 °C and 18,000 x *g*, 50 µL of the resulting mixture were used for quantification of creatinine using a Creatinine Assay Kit (ab204537, Abcam) as described in the manufacturer’s protocol. The remaining 50 µL was filtered through a Durapore PVDF 0.22-μm membrane using Ultrafree centrifugal filters (Millipore, UFC30GV00), and 5 µL was injected into the LC/MS.

For fecal pellets, ∼40 mg wet feces was pre-weighed into a 2 mL screw top tube containing six 6-mm ceramic beads (Precellys® CK28 Lysing Kit). 600 μL of a mixture of ice-cold acetonitrile, methanol, and water (4/4/2, v/v/v) were added to each tube and samples were homogenized by vigorous shaking using a QIAGEN Tissue Lyser II at 25/s for 10 min. The resulting homogenates were subjected to centrifugation for 15 min at 4 °C and 18,000 x *g*. 100 μL of the supernatant weas combined with 100 µL of internal standard water solution (20 μM 4-chloro-L-phenylalanine and 2 μM d^4^-cholic acid). The resulting mixtures were filtered through a Durapore PVDF 0.22-μm membrane using Ultrafree centrifugal filters (Millipore, UFC30GV00), or a MultiScreen Solvinert 96 Well Filter Plate (Millipore, MSRLN0410), and 5 µL was injected into the LC/MS.

#### Liquid chromatography/mass spectrometry (LC/MS)

For aromatic amino acid metabolites, analytes were separated using an Agilent 1290 Infinity II UPLC equipped with an ACQUITY UPLC BEH C18 column (1.7 μm, 2.1 mm x 150 mm, Waters Cat. #186002352 and #186003975) and detected using an Agilent 6530 Q-TOF equipped with a standard atmospheric-pressure chemical ionization (APCI) source or dual Agilent jet stream electrospray ionization (AJS-ESI) source operating under extended dynamic range (EDR 1700 *m*/*z*) in negative ionization mode. For the APCI source, the parameters were as follows: gas temperature, 350 °C; vaporizer, 350 °C; drying gas, 6.0 L/min; nebulizer, 60 psig; VCap, 3500 V; corona, 20 μA; and fragmentor, 135 V. For the AJS-ESI source, the parameters were as follows: gas temperature, 350 °C; drying gas, 10.0 L/min; nebulizer, 40 psig; sheath gas temperature, 300 °C; sheath gas flow, 11.0 L/min; VCap, 3500 V; nozzle voltage, 1400 V; and fragmentor, 130 V. Mobile phase A was H_2_O with 6.5 mM ammonium bicarbonate, and B was 95% MeOH with 6.5 mM ammonium bicarbonate. 5 µL of each sample was injected via autosampler into the mobile phase, and chromatographic separation was achieved at a flow rate of 0.35 mL/min with a 10 min gradient condition (*t*=0 min, 0.5% B; *t*=4 min, 70% B; *t*=4.5 min, 98% B; *t*=5.4 min, 98% B; *t*=5.6 min, 0.5% B).

For bile acids, compounds were separated using an Agilent 1290 Infinity II UPLC equipped with a Kinetex C18 column (1.7 µm, 2.1 x 100 mm, Phenomenex, Cat. #00D-4475-AN) and detected using an Agilent 6530 Q-TOF equipped with a dual Agilent jet stream electrospray ionization (AJS-ESI) source operating under extended dynamic range (EDR 1700 *m*/*z*) in negative ionization mode. The parameters of AJS-ESI source were as follows: gas temperature, 300 °C; drying gas, 7.0 L/min; nebulizer, 40 psig; sheath gas temp, 350 °C; sheath gas flow, 10.0 L/min; VCap, 3500 V; nozzle voltage, 1400 V; and fragmentor, 200 V. Mobile phase A was H_2_O with 0.05% formic acid, and B was acetone with 0.05% formic acid. 5 µL of each sample was injected via autosampler into the mobile phase and chromatographic separation was achieved at a flow rate of 0.35 mL/min with a 32 min gradient condition *(t=*0 min, 25% B; *t*=1 min, 25% B; *t*=25 min, 75% B, *t*=26 min, 100% B, *t*=30 min, 100% B, *t*=32 min, 25% B).

Online mass calibration was performed using a second ionization source and a constant flow (5 μL/min) of reference solution (119.0363 and 966.0007 *m*/*z*). The MassHunter Quantitative Analysis Software (Agilent, v. B.09.00) was used for peak integration based on retention time (tolerance of 0.2 min) and accurate *m*/*z* (tolerance of 30 ppm) of chemical standards. Quantification was based on 2-fold dilution series of chemical standards spanning 0.05 to 100 μM (aromatic amino acid metabolites) or 0.001 to 100 μM (bile acids) and measured amounts were normalized by weights of extracted tissue samples (pmol/mg wet tissue) or creatinine level in the urine sample (μM/mM creatinine). The linear quantification range and lower limit of detection for all metabolites are in **Table S5**. The MassHunter Qualitative Analysis Software (Agilent, version 7.0) was used for targeted feature extraction, allowing mass tolerances of 30 ppm.

#### *E. coli* colonization resistance

6-8-week-old female germ-free SW mice were orally gavaged with 200 µL of hCom1, hCom2, a stool sample from a healthy human donor, or 12Com—or with 220 µL of hCom2+Enteromix or 12Com+Enteromix—and fecal pellets were sampled weekly for 4 weeks. After 4 weeks, mice were orally gavaged with a 200 µL misture containing 10^9^ CFUs of EHEC and fecal pellets were sampled on days 0 (pre-EHEC infection), 2, 4, 6, and 14. After collection, all stool was prepared aerobically. Specifically, stool pellets were weighed and 10x w/v PBS was added to the tube. Each pellet was crushed with a 1 mL pipette tip and vortexed at maximum speed for 30 s to create a homogenous mixture. This mixture was serially diluted 1:10 six successive times and 5 µL of each dilution was plated on McConkey-Sorbitol agar. Plates were incubated at 37 °C for 16-18 h. The resulting colonies were enumerated and verified to be EHEC by metagenomic sequencing. Fecal pellets were also subjected to DNA extraction, metagenomic sequencing, and NinjaMap analysis to estimate strain relative abundances.

### QUANTIFICATION AND STATISTICAL ANALYSIS

Relative abundances were calculated from the output of NinjaMap or MIDAS without rarefying the total number of reads across samples. Relative abundances at each time point were averaged across the 4-5 mice that were co-housed in the same isolator and subjected to the same fecal challenge. Correlation coefficients were calculated after setting undetected bins to a minimum value (10^−6^ and 10^−8^ for MIDAS and NinjaMap, respectively) and performing a log_10_ transformation. Mice were not considered in the fecal challenge analyses if sequence reads in a sample from any week were of poor quality or abnormally variable. This affected one of five mice in all groups except for fecal challenge experiment 1, Hum3 (2 mice affected) and fecal challenge experiment 2, Hum1 (0 mice affected). Further details of statistical analyses can be found in the corresponding figure legends. All statistical analyses and tests were performed in MATLAB, and scripts for analyses are available at https://github.com/FischbachLab.

## ACKNOWLEDGMENTS

We are deeply indebted to members of the Fischbach and Huang labs for helpful discussions. This work was supported by a Dean’s Postdoctoral Fellowship (to P.-Y.H.), Human Frontier Science Program award LT000493/2018-L (K.N.), a Fellowship from the Astellas Foundation for Research on Metabolic Disorders (K.N.)., the Stanford Microbiome Therapies Initiative (M.A.F., K.C.H.), NIH grants DP1 DK113598 (M.A.F.), P01 HL147823 (M.A.F.), R01 DK101674 (M.A.F.), RM1 GM135102 (K.C.H.), and R01 AI147023 (K.C.H.), the Bill and Melinda Gates Foundation (M.A.F.), an HHMI-Simons Faculty Scholars Award (M.A.F.), the Leducq Foundation (M.A.F.), the Stanford-Coulter Translational Research Grants Program (M.A.F.), the Chan Zuckerberg Biohub (K.C.H., M.A.F.), and the Allen Discovery Center at Stanford on Systems Modeling of Infection (K.C.H.).

## AUTHOR CONTRIBUTIONS

Conceptualization: A.C., M.A.F. Methodology and investigation: A.C., P.-Y.H., S.J., X.M., M.W., F.B.Y., M.I., A.B., K.N., A.Z., A.P., K.A., A.W., R.Y., S.H., K.C.H., M.A.F. Formal analysis: A.C., P.-Y.H., S.J., X.M., M.W., M.I., K.C.H., M.A.F. Visualization: A.C., P.-Y.H., S.J., K.C.H., M.A.F. Supervision: N.N., J.L.S., K.C.H., M.A.F. Writing: A.C., P.-Y.H.., S.J., K.C.H., M.A.F. All authors reviewed the manuscript before submission.

