## Supplementary Information for "*In vivo* augmentation of a complex gut bacterial community"

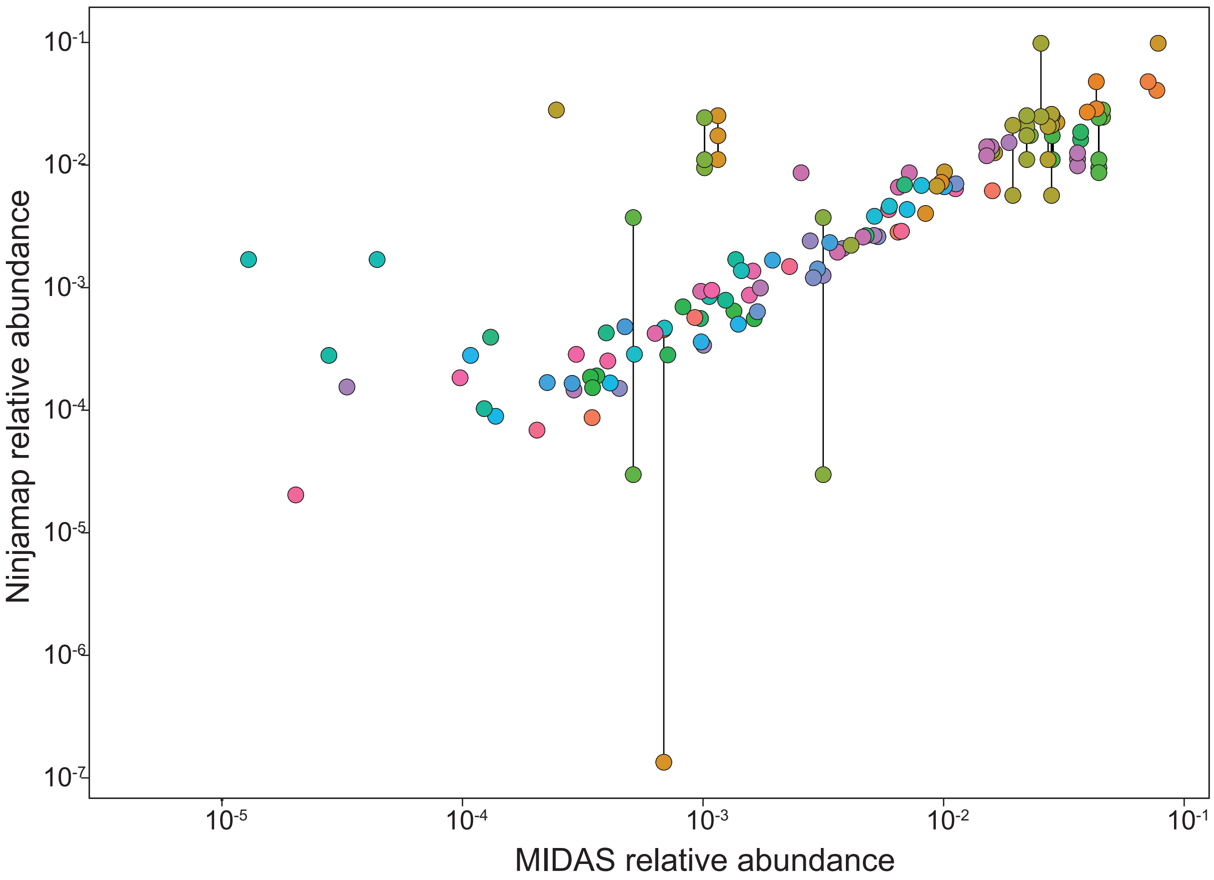


**Figure S1: Comparison of MIDAS and NinjaMap sequencing read utilization.** Mapping results are shown from the hCom2 inoculum used for the second fecal challenge experiment (**Figure 3**). Each dot represents a strain and the corresponding species bucket identified by MIDAS. Points connected by lines represent two strains that NinjaMap can distinguish but which map to the same MIDAS bucket, and are therefore colored similarly. Spearman's correlation coefficient is 0.83 and Pearson correlation coefficient is 0.717.

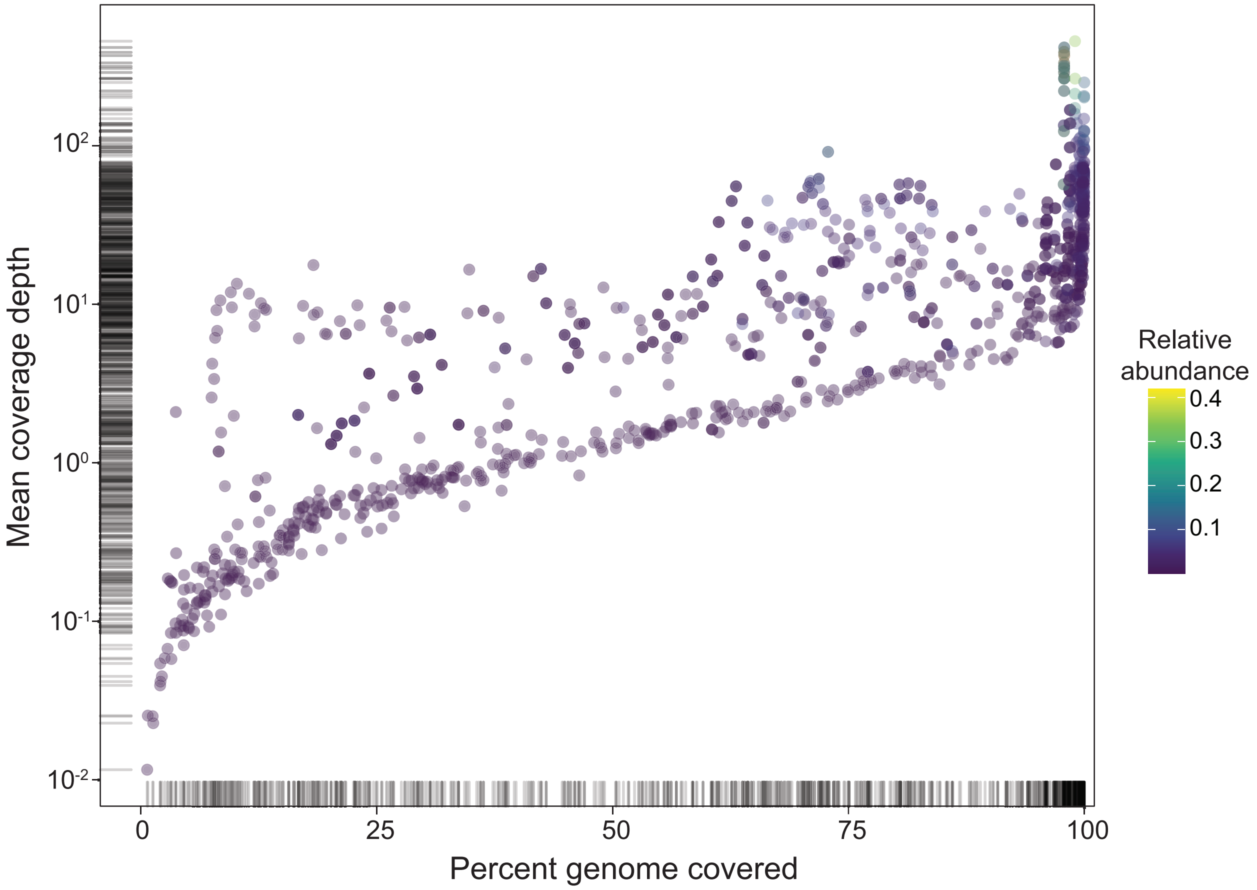


**Figure S2: NinjaMap strain coverage from the first fecal challenge experiment (Figure 2).** Reads from week 8 of the PBS challenge for hCom1 were aligned to the reference genomes in our database. Reads were retained only if they matched a database strain at 100% identity over 100% read length. Over 60% of strains had >95% breadth of coverage at 10x depth from perfectly matched reads.

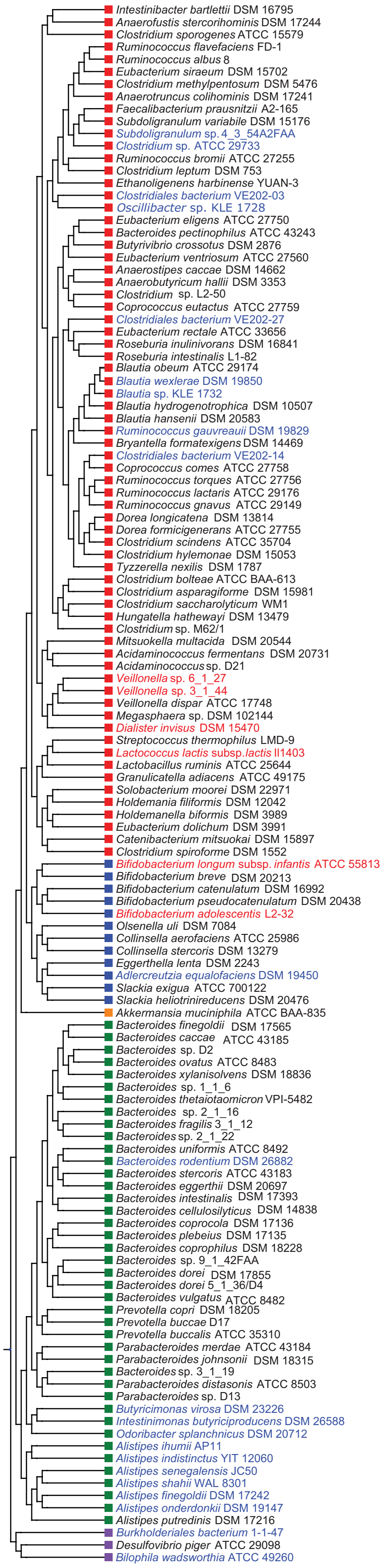


**Figure S3: Phylogenetic tree of strains from hCom1 and hCom2 strains.** Squares are colored according to phyla (red, Firmicute; blue, Actinobacteria; orange, Verrucomicrobia; green, Bacteroidetes; purple, Proteobacteria). Strain names in black are present in hCom1 and hCom2. Strain names in red were only in hCom1, whereas strain names in blue were only in hCom2.

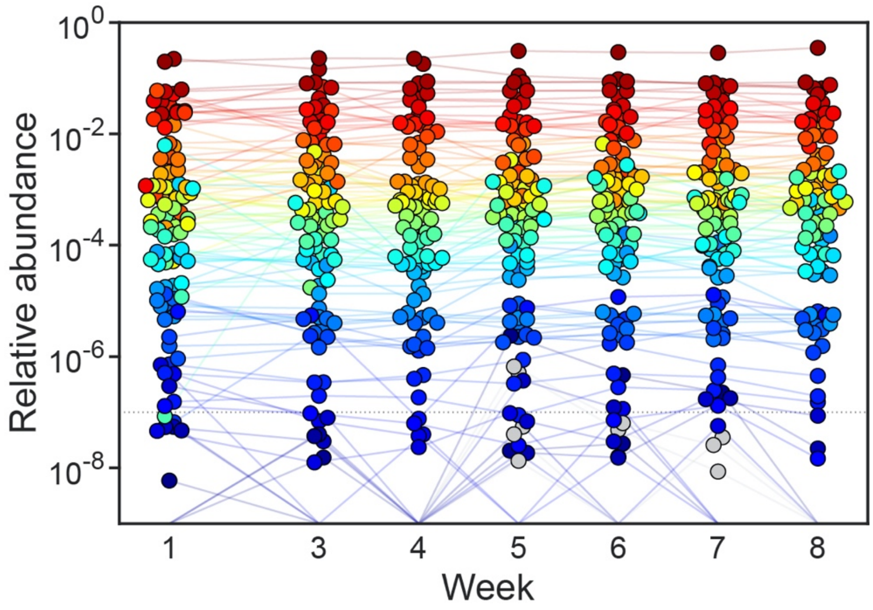


**Figure S4: hCom2 reaches a stable state soon after colonization.** Each dot is an individual strain; the collection of dots in a column represents the community at a single timepoint averaged over 5 mice co-housed in a cage. Strains are colored according to their rank-order relative abundance at week 4.

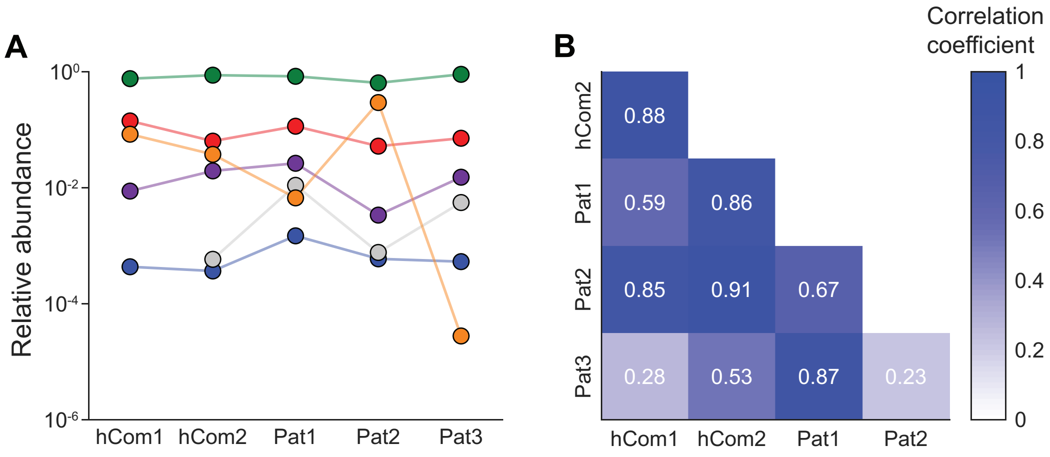


**Figure S5: The architecture of hCom2 more closely resembles that of a human fecal consortium than hCom1**. (**A**) Phylum-level relative abundances of hCom1, hCom2, and healthy human stool samples 1-3. **(B)** Pairwise correlation coefficients of relative abundance vectors were higher between hCom2 and the stool samples than hCom1 and the stool samples.

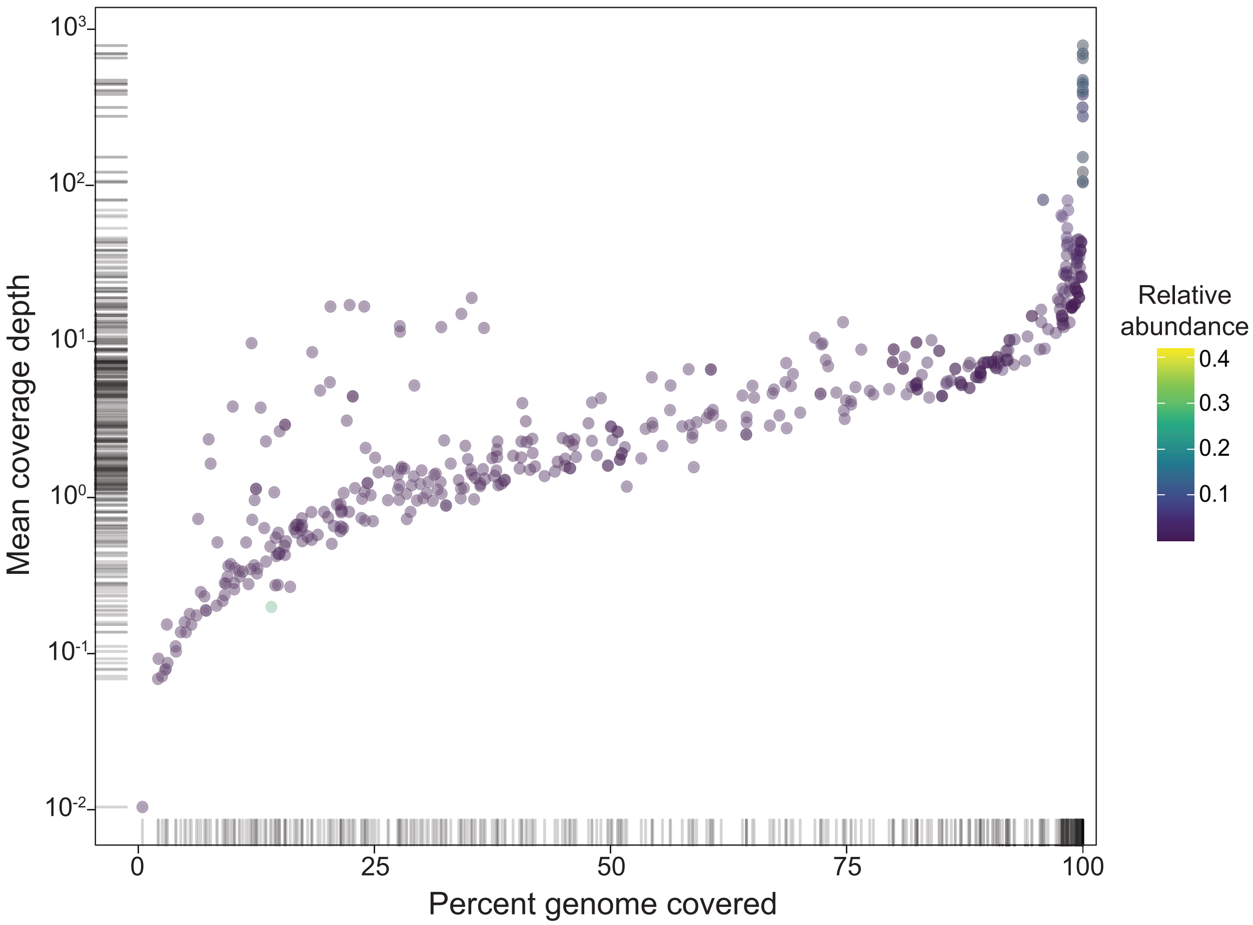


**Figure S6: NinjaMap strain coverage from the second fecal challenge experiment (Figure 3).** Reads from week 8 of the PBS challenge for hCom2 were aligned to the reference genomes in our databases. Reads were retained only if they matched a database strain at 100% identity over 100% read length. 54% of original strains in hCom2 and 76% of the new strains in hCom2 had >95% coverage of the of the reference genome by perfectly matched reads.

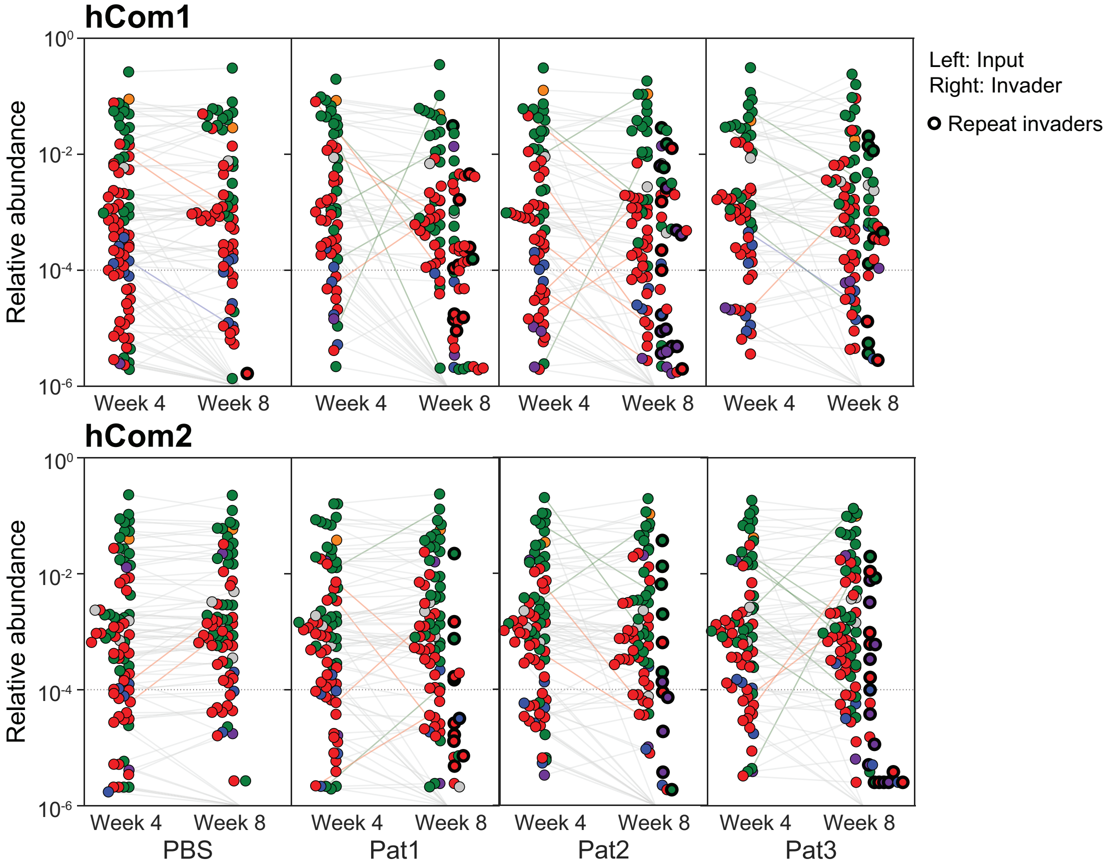


**Figure S7: Invading strains from the first (Figure 2) and second (Figure 3) fecal challenge experiments.** Averaged relative abundances of MIDAS bins—a rough proxy for species—are shown for hCom1- and hCom2-colonized mice. Week 4 and 8 species distributions are shown for each group. Week 8 distributions are split into two 'halves': the left half represents input species and the right half shows invading species. Invading species common to all three groups are outlined in bold; their total abundance significantly decreased from the first to the second fecal challenge experiment.

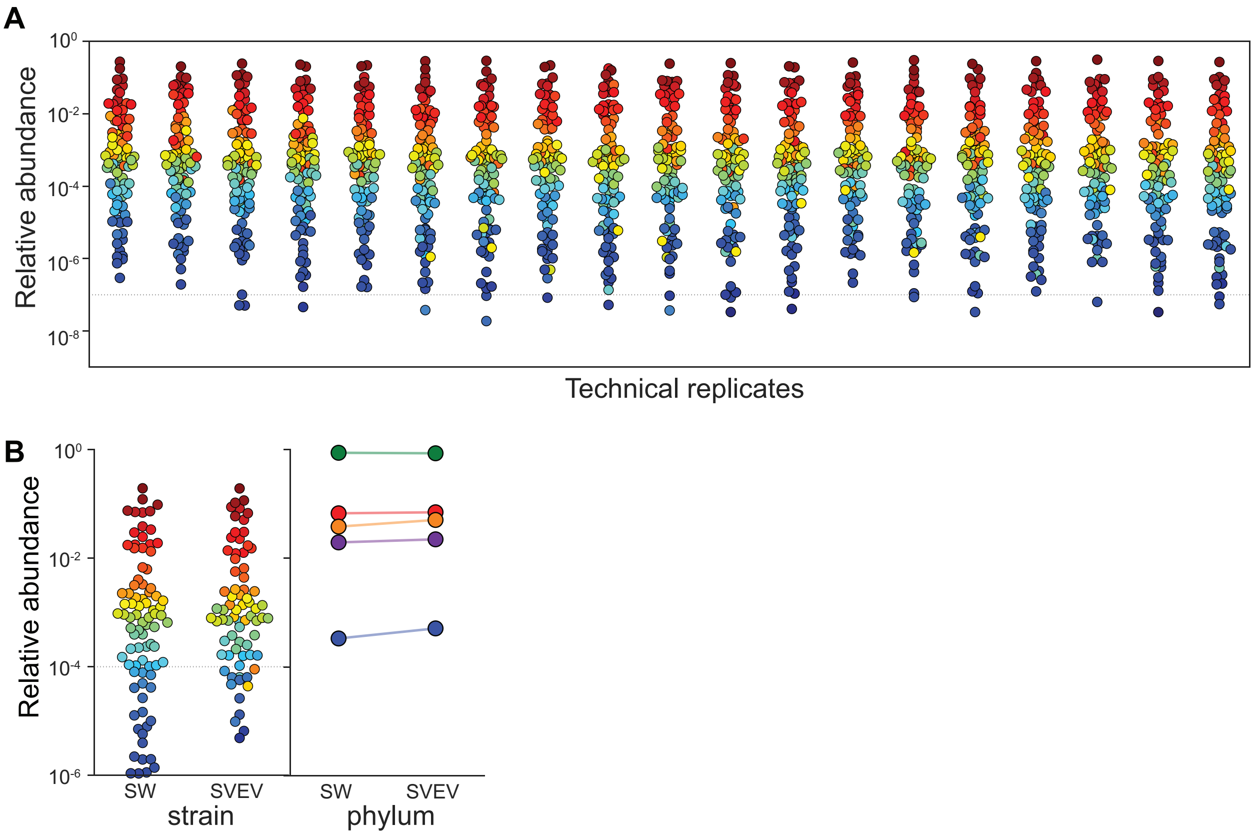


**Figure S8: The architecture of hCom2 is highly reproducible across technical replicates and similar between two strains of mice.** (**A**) Metagenomic sequencing of fecal samples from the second fecal challenge experiment (**Figure 3**)—prior to fecal challenge (Week 4)—reveals a high degree of technical reproducibility. As assessed by NinjaMap, strain relative abundances in the 20 mice colonized by the same hCom2 inoculum were highly similar (pairwise Pearson's correlation coefficients >0.93). **(B)** Two genetically distinct strains of mice, Swiss webster (SW) and SVEV, were colonized with hCom2 and housed for 4 weeks. Fecal community profiles are shown at the strain and phylum levels. Pearson correlation coefficient between SW and SVEV mice was >0.95.

### Supplemental Tables

**Table S1: Invading strains from the fecal challenge experiments.** Twenty-six bacterial species entered hCom1 from ≥2 of the 3 fecal samples used as a challenge. 42 species entered hCom1 from one of the 3 fecal samples. Strains that invaded or dropped out in each experiment are listed by sheet.

**Table S2: hCom2 strains and growth media**

**Table S3: hCom2 strain accession numbers**

**Table S4: MIDAS sensitivity analysis.**

**Table S5: Linear quantification range and lower limit of detection for bacterial metabolites in this study**.
